# Genomic insights into multidrug resistant *Escherichia coli* from bovine mastitis in Bangladesh

**DOI:** 10.1101/2025.10.21.683615

**Authors:** Naim Siddique, Kh. Yeashir Arafat, Md Abu Ahsan Gilman, Md. Morshedur Rahman, Ziban Chandra Das, Tofazzal Islam, M. Nazmul Hoque

## Abstract

**Background:** Mastitis poses a significant threat to dairy industry and public health due to the emergence of multidrug-resistant (MDR) *Escherichia coli*. This study provides a genomic characterization of two MDR *E. coli* strains, MBBL4 and MBBL5, from bovine mastitis in Bangladesh, highlighting their evolutionary relationships, resistome, and virulome.

**Methods:** Species-level identification of MBBL4 and MBBL5 was confirmed using biochemical assays, VITEK-2 system, and *16S rRNA* gene sequencing. Antimicrobial susceptibility profiling was conducted to determine their resistance patterns. Whole genome sequencing (WGS) and comprehensive genomic analysis were performed for phylogenetic, comparative genomics, mobile genetic elements (MGEs), antimicrobial resistance genes (ARGs), and virulence factor genes (VFGs) analyses.

**Results:** Both isolates exhibited extensive MDR patterns, showing resistance to ten antibiotics. Phylogenetic and ANI analyses showed that MBBL4 clustered with mastitis-associated and human bacteremia strains of *E. coli*, while MBBL5 was closely related to wildlife-associated strains, reflecting divergent evolutionary lineages. Pangenome analysis revealed an open pangenome structure, indicating high genetic diversity, with MBBL4 harboring 21 unique genes and MBBL5 possessing nine unique genes. Both genomes harbored numerous ARGs spanning over 11 antibiotic classes, and VFGs, predominantly associated with adherence and secretion systems, underscoring their extensive resistome, virulome, and adaptive potentials. Abundant MGEs (plasmids, prophages, insertion sequence elements and genomic islands) further underscored the role of horizontal gene transfer in driving resistance and virulence in these strains.

**Conclusion:** This study highlights the zoonotic potential and adaptive capacity of MDR *E. coli* from bovine mastitis in Bangladesh driven by resistome, virulome, and mobile genetic elements. These findings highlight the urgent need for One Health-based genomic surveillance to mitigate MDR *E. coli* transmission from dairy farms to humans and the environment.

## Background

Mastitis is an inflammatory condition that leads to pathological changes in the mammary glandular tissue [1]. It incurs substantial economic losses in dairy industry by adversely affecting animal health, compromising milk quality, and leading to substantial financial losses for dairy farmers [2]. Various etiological agents are involved in the occurrence of mastitis. Among them, bacteria are the most common etiological agents, followed by fungal, viral and mycoplasma infections [3]. Contagious and environmental bacterial pathogens are responsible for causing majority of the mastitis infections [4]. Among the environmental bacterial pathogens implicated in mammary infections, *E. coli* is the most prevalent, followed by species of *Klebsiella* and *Enterobacter* spp. [5, 6]. *E. coli* is a prominent environmental pathogen associated with both acute and subclinical forms of mastitis [7, 8]. Beyond its impact on animal health and milk production, *E. coli* from bovine sources is increasingly recognized as a potential vector for AMR genes [9, 10]. The increasing prevalence of AMR in livestock, particularly the dissemination of antimicrobial-resistant *E. coli* strains, poses a significant zoonotic and public health concern [11, 12]. The global rise of antimicrobial resistance among *E. coli* strains complicates the treatment of mastitis and represents a broader threat to both veterinary and human medicine [13, 14]. In addition, *E. coli* is an important human pathogen causing urinary tract infections, neonatal meningitis, and septicemia, with ESBL-producing strains posing a serious One Health concern due to resistance to critical β-lactam antibiotics [15].

Antimicrobial-resistant *E. coli* commonly harbor ARGs and VFGs, which are key contributors to drug resistance and infection development [16]. Their ability to disseminate across hosts and transfer these genes facilitates the accumulation of additional resistance traits, driving the emergence of MDR *E. coli* strains [17]. Mobile genetic elements (MGEs) in bacteria are DNA segments that can move within or between bacterial cells, facilitating horizontal gene transfer (HGT) and driving bacterial evolution and adaptation [18]. Among MGEs, plasmids are particularly significant, acting as vehicles for the transmission of ARGs, VFGs, and other functional genes. They are classified as conjugative or mobilizable based on their ability to mediate gene transfer, although not all plasmids are self-transmissible [19]. The detection of MGEs and regions of genomic plasticity (RGPs) in pathogenic *E. coli* reflects extensive genomic evolution largely driven by HGT, which enables the rapid acquisition of complex traits such as AMR and virulence [20]. Furthermore, the detection of distinct genomic islands (GIs) in *E. coli* isolates often carrying additional ARGs, VFGs, and mobile elements further supports the notion that MGEs play a pivotal role in the evolution and dissemination of pathogenic traits [21, 22]. These observations highlight the critical need for ongoing surveillance of *E. coli* associated with bovine mastitis, as resistance determinants from these strains have the potential to transfer to human pathogens. Therefore, a comprehensive understanding of the genomic architecture of bovine mastitis-associated *E. coli* is essential for uncovering resistance mechanisms, monitoring evolutionary dynamics, and informing effective surveillance and mitigation strategies.

Genomic approaches, particularly WGS, now provide powerful tools to explore these mechanisms at high resolution and to track the evolutionary pathways of antimicrobial resistance and virulence in mastitis-associated pathogens within a One Health framework [23, 24]. Similar applications have been demonstrated in recent genomic risk assessments of *Salmonella enterica* from food animals in China, highlighting the effectiveness of WGS-based surveillance across the food chain [25]. Gazipur, one of the dairy hubs in Bangladesh where this study was conducted, has limited genomic data on mastitis related *E. coli*, providing little insight into the genomic determinants of AMR, virulence, and HGT in local isolates from poultry, wastewater, and environmental sources [26–28]. Epidemiological studies report a pooled prevalence of 39.05% for subclinical mastitis and 11.18% for clinical mastitis (CM) in dairy cows in Bangladesh [29], with a substantial proportion of mastitis pathogens exhibiting MDR phenomena [30–32] largely attributable to the irrational and excessive use of antibiotics. These findings underscore the urgent need for continued surveillance and evidence-based antimicrobial management to limit the spread of MDR pathogens. Given this background, we performed high-throughput genome sequencing of two MDR *E. coli* isolates, MBBL4 and MBBL5, screened from bovine CM milk samples collected from two small-holding dairy farms in Gazipur district of Bangladesh. This study offers an in-depth genomic characterization of these mastitis-associated *E. coli* strains, revealing the significant roles of MGEs, ARGs, VFGs, and genomic plasticity in their evolution. The results underscore the pressing need for ongoing animal-human AMR surveillance to track potential cross-species transmission and mitigate the spread of MDR bacterial clones.

## Methods

### Sample collection and isolation of bacteria

Milk samples (n = 45) were aseptically collected from 45 lactating cows showing clinical signs of mastitis, located across 30 smallholder dairy farms (each housing 5–20 cows) in the Gazipur district of Bangladesh (24.09° N, 90.41° E) between January 2024 and May 2025. Samples were collected, and prevalence was calculated based on observed frequencies without applying any statistical procedures. CM was confirmed in all selected cows based on observable clinical signs, such as changes in milk color, udder swelling, redness, and elevated temperature, along with a California Mastitis Test (CMT) score of 3, indicating the presence of numerous clots [33]. During the morning milking session (8:00–10:00 a.m.) approximately 10 mL of milk was aseptically collected from each cow into sterile Falcon tubes after careful disinfection of the teat ends. Strict adherence to hygiene protocols was maintained throughout the process to minimize contamination and ensure sample integrity. For the isolation and phenotypic identification of *E. coli*, samples were first enriched in nutrient broth (Oxoid, UK) with overnight incubation at 37°C. The enriched broth was then serially diluted up to 10^−3^ fold, and a portion (2 μl) was plated onto Eosin Methylene Blue (EMB) agar (Oxoid, UK) and incubated at 37°C for 24 hours. Two to three suspected colonies exhibiting a characteristic green metallic sheen were selected per sample and sub-cultured for re-isolation. Presumptive *E. coli* isolates were identified based on Gram staining (gram-negative short rods) and a series of biochemical tests including catalase, oxidase, urease, indole, methyl red, Voges-Proskauer (VP), and triple sugar iron (TSI) tests [14, 31]. Species-level identification was subsequently confirmed using the VITEK-2 system v9.01 [34] and ribosomal gene (*16S rRNA*) sequencing using UNI8f 5′-AGAGTTTGATCCTGGCTCAG-3′ and uni 1492r 5′-GGTTACCTTGTTACGACTT-3′ primers [31].

### Antimicrobial susceptibility profiling

The antimicrobial susceptibility testing (AST) profiles of *E. coli* isolates (n = 33) were assessed using the Kirby-Bauer disc diffusion method [32], in accordance with the Clinical Laboratory Standards Institute (CLSI) M100, 34th edition guidelines [35]. The isolates were tested against a panel of 15 antibiotics widely used for treating bacterial infections in both humans and animals, including bovine mastitis in Bangladesh, with AST performed without replication. The antibiotics tested were Beta-lactams (ampicillin, 10 μg/ disc; oxacillin, 1 μg/ disc), Monobactams (aztreonam, 30 μg/ disc), Tetracyclines (doxycycline, 30 μg/ disc; tetracycline, 30 μg/ disc), Nitrofurans (nitrofurantoin, 300 μg/ disc), Fluoroquinolones (ciprofloxacin, 10 μg/ disc; nalidixic acid, 30 μg/ disc), Cephalosporins (cefoxitin, 30 μg/ disc), Carbapenems (imipenem, 10 μg/ disc), Aminoglycosides (gentamycin, 10 μg/ disc; streptomycin, 10 μg/ disc), Chloramphenicol (30 μg/ disc), Macrolides (azithromycin, 15 μg/ disc) and Sulphonamides (sulphonamide compound, 300 μg/ disc). Resistance break-point was interpreted using CLSI guidelines. *E. coli* isolates (n = 27) demonstrating resistance to ≥3 antibiotic classes were classified as MDR.

### Genome sequencing, assembly, and annotation

Genomic DNA from two highly resistant MDR *E. coli* isolates (MBBL4 and MBBL5), exhibiting resistance to 10 antibiotics, was extracted following established standard protocols [36, 37]. In brief, high-quality genomic DNA from *E. coli* isolates MBBL4 and MBBL5 was extracted from nutrient broth cultures using the QIAamp DNA Mini Kit (QIAGEN, Germany) following the manufacturer’s protocol. DNA libraries were prepared from 1 ng of input DNA utilizing the Nextera DNA Flex Library Preparation Kit (Illumina, USA). Thereafter, WGS was subsequently performed on the Illumina MiSeq platform employing a 2 × 250 bp paired-end sequencing run [14, 30]. The quality of raw WGS reads was evaluated using FastQC v0.11.7 [38].Trimmomatic v.0.39 [39] was used for trimming Illumina adapters, known Illumina artifacts, phiX reads. Subsequently, genome assembly was done with SPAdes v.3.15.5 [40] and annotated using the NCBI Prokaryotic Genome Annotation Pipeline v.6.6 [41]. Genome completeness was also estimated using CheckM v1.2.3 [42] analysis. The RAST v.2.0 [43] web-server was employed to count the number of subsystems, while PathogenFinder v.1.1 [44] was used to predict the probability of the isolated strains as human pathogens. *E.coli* strains MBBL4 and MBBL5 underwent genomic visualization and functional annotation using Genovi v0.2.16 [45]. The number of genes that are included to each Clusters of Orthologous group (COG) functional category was predicted and visualized via the bar graphs. All software were used with default parameters unless specified.

### Comparative phylogenetic analysis

For phylogenetic analysis, 30 *E. coli* genomes were used, including our two isolated genomes (MBBL4 and MBBL5) and 28 reference genomes from NCBI (**Table S1**). These genomes were sourced from diverse origins, with 17 from bovine, six from human, five from other animals, and one each from water and soil. The Mashtree v2.0 [46] was used to generate a mash distance-based neighbour-joining phylogenetic tree to assess genetic relatedness and their evolutionary relationships. The tree was illustrated with iTOL v7 [47]. All 30 *E. coli* genomes were subjected to average nucleotide identity (ANI) analysis using ANIclustermap v.1.3.0 [48] and pangenome analysis using Roary v3.12.0 [49]. The pangenome was categorized into four gene classes based on their prevalence across strains: core genes (present in 99–100% of strains), soft core genes (95–<99%), shell genes (15–<95%), and cloud genes (0–<15%). Additionally, pangenome and core genome curves were generated for 17 closely related *E. coli* genomes including MBBL4 and MBBL5 (based on phylogenetic analysis), using the PanGP v1.0.1 tool [50]. New gene discovery was modeled as a function of the sequential addition of genomes, employing the distance guide algorithm with 100 replicates and 3,000 genome order permutations. PanGP applies a power-law regression model (*y = Ax^B^ + C*) to estimate the pangenome size, where *y* represents the total number of gene families, *x* is the number of genomes, and *A*, *B*, and *C* are fitting parameters. When 0 < B < 1, the pangenome should be considered an open pangenome. For core genome estimation, an exponential decay model (*y = Ae^Bx^ + C*) was applied, where *y* is the core genome size and *x* is the number of genomes. Similarly, new gene discovery was fitted using a least-squares power-law model (*y = Ax^B^*), where *y* represents the number of newly identified genes and *x* corresponds to the number of genomes, with *A* and *B* as fitting parameters [51].

### Genome plasticity analysis

The 17 *E. coli* genomes, including MBBL4 and MBBL5, were analyzed for mobile genomic elements (MGEs). We used panRGP [52] to detect the regions of genomic plasticity (RGPs), and ISseeker [53] to identify insertion sequences (ISs). Prophage regions and plasmids were identified by PHASTEST [54] and Platon [55], respectively.

### Identification of antibiotic resistance genes, virulence determinants, and genomic islands

Antibiotic resistance genes (ARGs) of 17 closely related *E. coli* genomes, including MBBL4 and MBBL5, were predicted via CARD v4.0.1 [56],while virulence factor genes (VFGs) were predicted by VFDB v6.0 [57] and ecoli_VF in ABRicate (https://github.com/tseemann/abricate) tool with default parameter, applying a minimum coverage and identity threshold of >80%. Additionally, Islandviewer4 [58] was used to predict genomic island of these selected genomes.

## Results

### Prevalence and phenotypic resistance profile of *E. coli* isolates

Thirty-three (n = 33) isolates of *E. coli* were screened from 45 samples using conventional culture techniques and Gram staining. Species-level identification of *E. coli* was confirmed through the VITEK-2 system and *16S rRNA*-gene sequencing. The overall prevalence of *E. coli* in bovine CM milk samples from the Gazipur district of Bangladesh, was 73.33%. AST profiling revealed that 81.82% (27/33) of the isolates were MDR, exhibiting resistance to ≥3 antibiotics. Among the 33 *E. coli* isolates, the highest resistance was observed against tetracycline (97.2%), followed by doxycycline (92.5%), sulfonamides (87.2%), nitrofurantoin (80.5%), and ciprofloxacin (78.6%). Additionally, these isolates showed 50–75% resistance to chloramphenicol, ampicillin, oxacillin, streptomycin, and gentamycin. Notably, isolates MBBL4 and MBBL5 exhibited extensive resistance profile, showing resistance to 10 antibiotics. The high resistance rates to tetracycline, doxycycline, and sulfonamides, commonly used for treatment of infectious diseases in Bangladesh, suggest that standard therapeutic regimens may be ineffective, emphasizing the need for informed antibiotic selection.

### Genomic features of *E. coli* strains MBBL4 and MBBL5

The assembled genomes of *E. coli* MBBL4 (NCBI accession no: JBINJR000000000) and MBBL5 (NCBI accession no: JBINJQ000000000) were approximately 4.6 Mbp in size, each with a GC content of 57.5%. The assemblies comprised 125 contigs for MBBL4 and 71 contigs for MBBL5. A total of 4,398 coding sequences (CDSs) were identified in MBBL4 and 4,385 in MBBL5, with 4,196 and 4,277 annotated as protein-coding genes, respectively. Quality assessment using CheckM demonstrated high completeness for both genomes, with 98.53% for MBBL4 and 99.56% for MBBL5, alongside minimal contamination levels of 0.27% and 0.70%, respectively. Furthermore, comprehensive genomic features including RNA genes, CRISPR-Cas arrays, prophage elements, ARGs, VFGs, and functional subsystem classifications identified in MBBL4 and MBBL5 genomes are detailed in **Table 1**. Both strains were predicted to be human pathogens, with pathogenicity scores of 0.857 (MBBL4) and 0.887 (MBBL5), exceeding the >0.80 high-risk threshold and corresponding to 136 and 155 pathogenic gene families, respectively. The circular genome representations of *E. coli* strains MBBL4 and MBBL5, along with their COG-based functional annotations, are illustrated in **Figs. 1A** and **1B**, respectively. Functional annotation assigned 4,678 genes in MBBL4 and 4,781 genes in MBBL5 into 25 COG functional categories distributed across four major functional groups. The most abundant COG category in both genomes was G (carbohydrate transport and metabolism), comprising 9.21% in MBBL4 and 9.30% in MBBL5, followed by category E (amino acid transport and metabolism), accounting for 8.23% and 8.26% in MBBL4 and MBBL5, respectively.

**Fig. 1.**
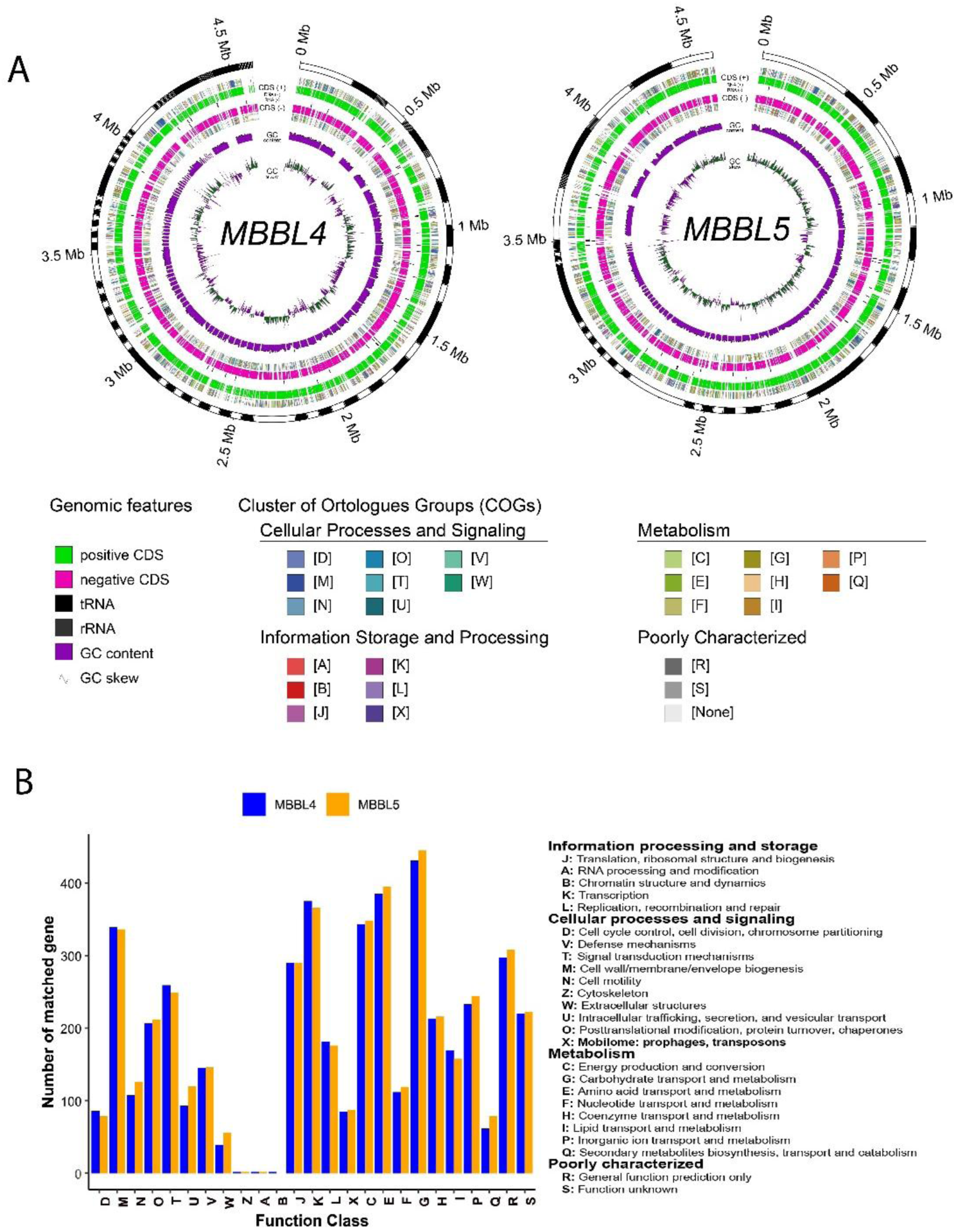
Genome visualization and functional annotation of *E. coli* strains MBBL4 and MBBL5. **(A)** Circular genome maps of MBBL4 and MBBL5. From the outermost to the innermost rings: contigs, coding sequences (CDS) on the forward strand with clusters of orthologous group (COG) category annotations, tRNA and rRNA genes, CDS on the reverse strand with COG category annotations, GC content, and GC skew. Different colors represent various features, as indicated in the legend. **(B)** Distribution of functional COG categories in the MBBL4 and MBBL5 genomes. Blue bars represent MBBL4, and orange bars represent MBBL5.

**Table 1.** Genomic features of *E. coli* strains MBBL4 and MBBL5.

| Genomic features | *E. coli* strains |  |
| --- | --- | --- |
|  | MBBL4 | MBBL5 |
| GenBank accession | <u>JBINJR000000000</u> | <u>JBINJQ000000000</u> |
| BioSample accession | <u>SAMN44249571</u> | <u>SAMN44249572</u> |
| SRA accession | <u>SRR30951065</u> | <u>SRR30951064</u> |
| Assembled genome size (bp) | 4,561,349 | 4,600,128 |
| GC % | 50.5 | 51 |
| Coverage (x) | 100 | 100 |
| Genome completeness (%) | 98.53 (0.27% contamination) | 99.56 (0.7% contamination) |
| N<sub>50</sub> value (bp) | 110283 | 194709 |
| Number of contigs | 125 | 71 |
| Genes (total) | 4,469 | 4,448 |
| CDSs (total) | 4,398 | 4,385 |
| Genes (coding) | 4,196 | 4,277 |
| CDSs (with protein) | 4,196 | 4,277 |
| Genes (RNA) | 71 | 63 |
| rRNAs | 2, 1, 2 (5S, 16S, 23S) | 2, 1, 1 (5S, 16S, 23S) |
| tRNAs | 54 | 52 |
| ncRNAs | 12 | 7 |
| Pseudo genes (total) | 202 | 108 |
| CRISPR arrays | 2 | 2 |
| Number of ARGs | 52 | 45 |
| Number of VFGs | 126 | 185; |
| Number of subsystems | 376 | 376 |
| Pathogenecity score (out of 1) | 0.857 | 0. 887 |
| Matched with pathogenic families | 136 | 155 |

(Note: N<sub>50</sub> is the length at which 50% of the genome is contained in contigs of this size or longer;
Coverage indicates the average number of times each base was sequenced).

### Genomic diversity and strain-specific genes in *E. coli* MBBL4 and MBBL5

Phylogenetic analysis revealed that *E. coli* MBBL4 and MBBL5 clustered into distinct clades, indicating divergent evolutionary lineages (**Fig. 2A**). Strain MBBL4 clustered with *E. coli* strains 1303 and D6-117_07.11, both previously isolated from bovine milk samples associated with mastitis. Additionally, MBBL4 exhibited notable genetic similarity to the human *E. coli* strain 5264, which was recovered from a case of human bacteremia. In contrast, MBBL5 was phylogenetically closer to *E. coli* strains A2 and A17, both fecal isolates obtained from wildlife species, the platypus and otter, respectively. ANI analysis, using a 95% nucleotide identity threshold for species delineation [59], further corroborated these phylogenetic relationships, demonstrating high genomic similarity between MBBL4, MBBL5, and their respective closest reference genomes (**Fig. 2B**). ANI cluster map analysis demonstrated that both MBBL4 and MBBL5 shared over 96% nucleotide identity with all reference *E. coli* genomes. Notably, MBBL4 and MBBL5 exhibited 98.5% nucleotide similarity to each other, indicating a close genetic relationship despite distinct phylogenetic clustering. Specifically, MBBL4 showed the highest ANI value of 99.3% with *E. coli* strains 21415616, P4, VL2874, VL2732, and 01T-32_03. In contrast, MBBL5 displayed the highest ANI similarity, approximately 99.7%, with *E. coli* strains A2, RHB33-C23, and D217-5 (**Fig. 2B**).

**Fig. 2.**
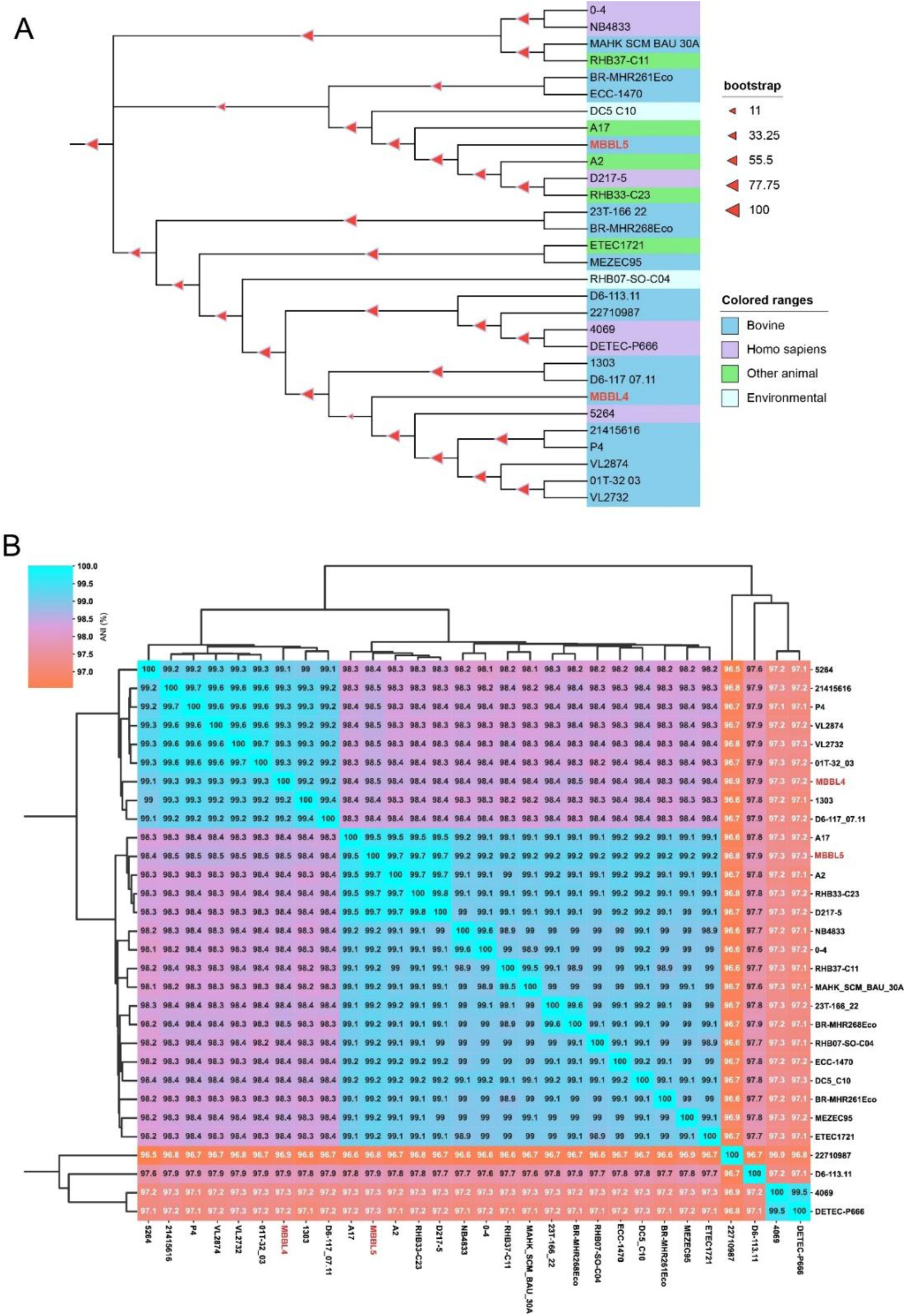
Overview of the evolutionary dynamics and average nucleotide identity (ANI) of *E. coli* strains MBBL4 and MBBL5 compared with 28 reference *E. coli* genomes. **(A)** Mash-distance-based neighbor-joining phylogenetic tree of 30 E. coli genomes. The sources of the isolates are indicated using distinct color codes. **(B)** Heatmap of ANI values displaying the clustering patterns among the 30 E. coli genomes.

A pangenome analysis was also conducted on our study genomes MBBL4 and MBBL5 along with reference genomes (**Table S1**) to elucidate their gene repertoire. The resulting pangenome dendrogram revealed that *E. coli* MBBL4 clustered closely with eight strains such as *E. coli* D6-117.07, 1303, 5264, P4, 21415616, VL2732, VL2874, and 01T-32/03 while MBBL5 showed the highest similarity to strains A2, RHB33-C23, and D217-5 (**Fig. 3A**). The pangenome analysis identified 13,571 genes across 30 *E. coli* genomes, comprising 2,983 core genes, 311 soft-core genes, 2,003 shell genes, and 8,268 cloud genes (**Fig. 3B**). Additionally, we identified 2,883 core genes in 17 closely related *E. coli* genomes through pangenome analysis (**Table S2**). The pangenome analysis demonstrated an open pangenome model (**Fig. 3C**), with the total gene cluster size continuously increasing as more genomes were added (y = 1131.59x^0.17^ + 2242.15; R² = 0.9993), indicating substantial genetic diversity. Conversely, the core genome exhibited a declining trend, fitting an exponential decay model (y = 540.66e^−0.24x^ + 2901.45; R² = 0.9724), suggesting gene loss with additional genomes. The new gene curve (**Fig. 3D**) followed a power-law decay (y = 247.28x^−0.86^; R² = 0.9753), reflecting that the number of novel genes decreases progressively as the genome number increases but does not plateau, supporting an open genome concept. Furthermore, strain-specific analysis revealed that MBBL4 harbored 21 unique genes and 380 accessory genes, whereas MBBL5 possessed 9 unique genes and 468 accessory genes (**Fig. 3E**, **Table S2**).

**Fig. 3.**
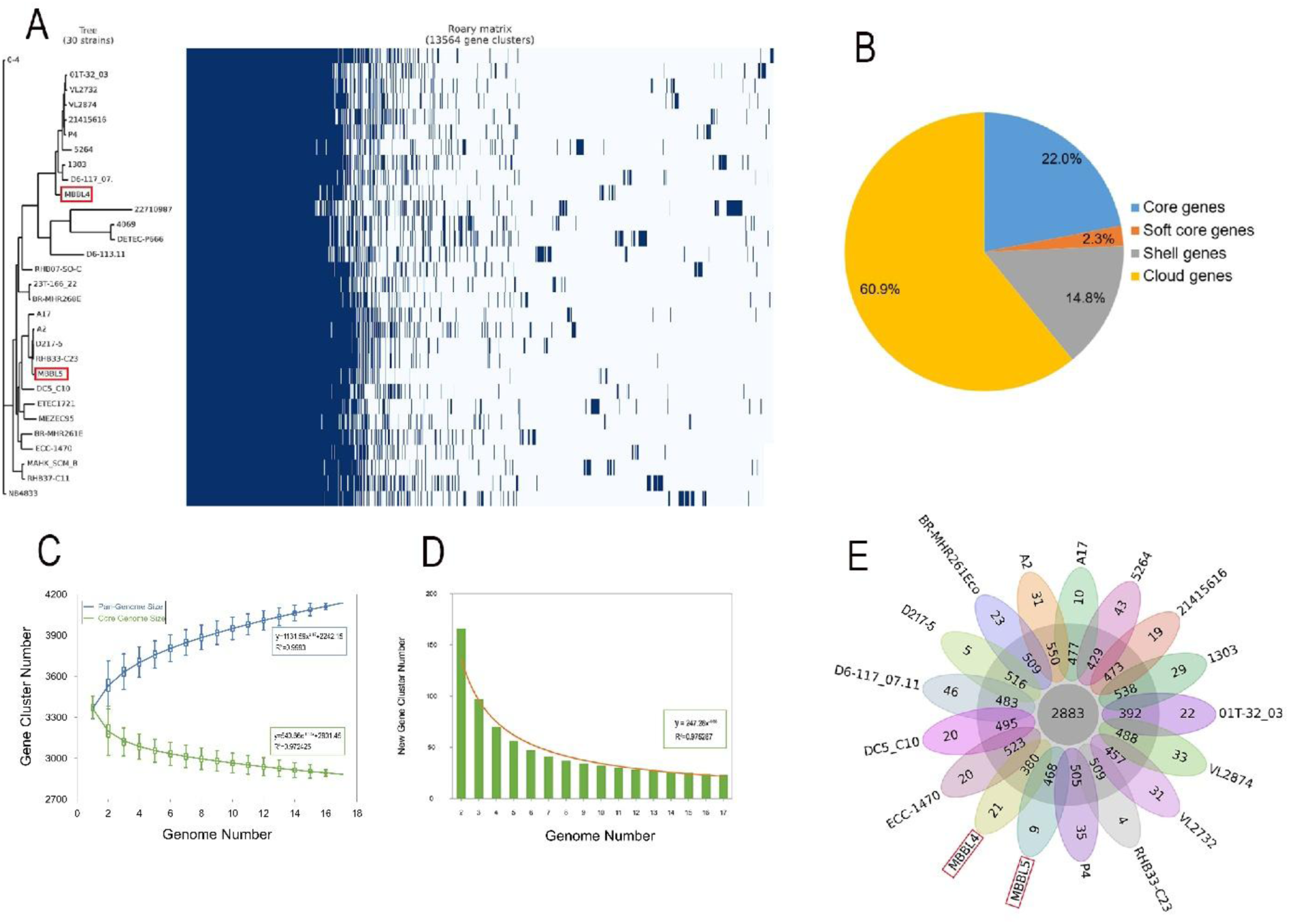
Overview of pangenome analysis of *E. coli* strains MBBL4 and MBBL5 compared with other reference *E. coli* genomes. **(A)** Gene presence/absence matrix based on pangenome clustering of MBBL4 and MBBL5 (highlighted in red) alongside 30 closely related *E. coli* genomes. **(B)** Distribution of gene frequencies across the 30 *E. coli* genomes. **(C)** Gene accumulation curves showing pangenome expansion (blue) and core genome reduction (green) with increasing numbers of genomes. Blue and green boxes indicate the sizes of the pangenome and core genome, respectively, at each step. **(D)** Bar graph representing the number of newly identified genes as additional genomes are incorporated into the analysis. **(E)** Flower plot illustrates the gene clusters across 17 *E. coli* genomes. The central circle indicates the conserved core genome shared by all isolates, surrounded by a ring of accessory genome containing partially shared genes and each petal representing the unique genes specific to a single strain, with petal length proportional to gene count. Blue boxes highlight MBBL4 and MBBL5 strains which exhibit genomic divergence from other isolates.

### Genomic plasticity and mobile genetic elements in *E. coli* MBBL4 and MBBL5

Genomic plasticity analysis of 17 phylogenetically related *E. coli* genomes revealed considerable variation in the number of RGPs (528), including 29 in MBBL4 and 21 in MBBL5 (**Table S3**). Furthermore, major MGEs, including ISs, prophages, and plasmids, were comprehensively analyzed as key drivers of genomic plasticity and evolutionary adaptation in these genomes (**Fig. 4**). A total of 773 IS elements, representing 19 distinct IS families, were identified across the 17 *E. coli* genomes (**Table S4**). The IS3 family was the most prevalent, with 176 elements, followed by the IS1 family with 143 elements. The number of IS elements varied significantly between the two strains, with MBBL5 possessing 14 and MBBL4 containing 29 (**Table S4**). Notably, several IS elements detected in the MBBL4 and MBBL5 genomes, including IS1, IS6, IS21, and IS30, may be linked to antimicrobial resistance. Prophage analysis revealed four prophage regions in *E. coli* MBBL4 compared to only two in MBBL5 (**Fig. 4, Table S5**). Plasmid analysis further differentiated the strains, identifying a 39.27 kb circular plasmid (p0111; GenBank accession no. AP010962) in MBBL4, while no plasmids were detected in MBBL5 (**Fig. 4** and **Table S6**).

**Fig. 4.**
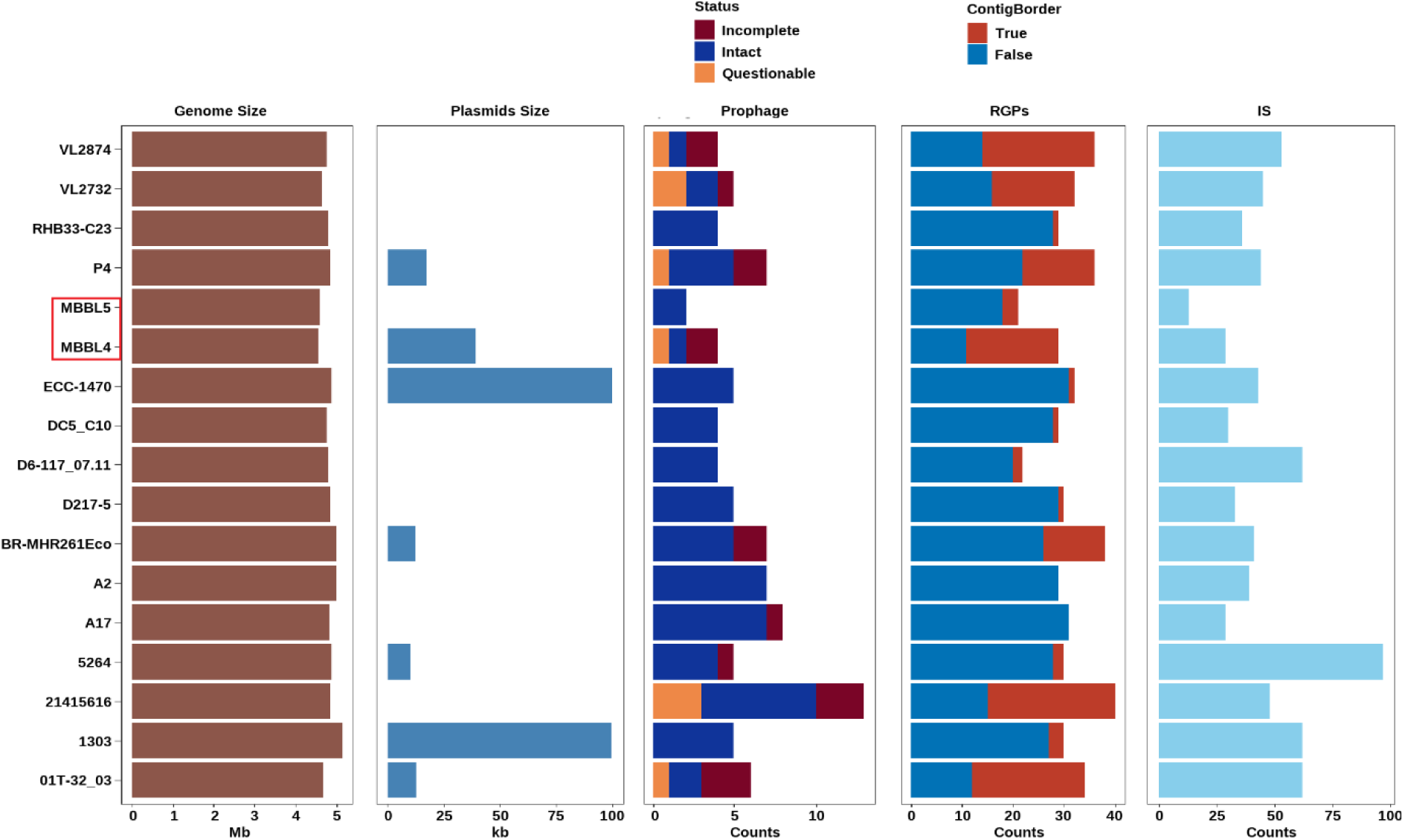
Overview of regions of genomic plasticity (RGPs) and mobile genetic elements (MGEs) identified in *E. coli* strains MBBL4, MBBL5, and 15 other closely related *E. coli* genomes. The figure displays, from left to right, the genome size, total plasmid size, number of prophages, number of RGPs, and number of insertion sequence (IS) elements for each *E. coli* genome. Both MBBL4 and MBBL5 strains are highlighted in red box.

### Resistome repertoire in *E. coli* MBBL4 and MBBL5

Comprehensive resistome analysis across 17 closely related *E. coli* genomes identified 128 ARGs distributed across 11 antibiotic classes. Of these, 59 ARGs were detected in the mastitis-associated strains MBBL4 and MBBL5, with 52 presents in MBBL4 and 45 in MBBL5 (**Fig. 5**). Importantly, 25 ARGs were found to be conserved in genomes (MBBL4 and MBBL5), showing resistance to aminoglycosides (*e.g., TolC*, *acrD, baeR*, *kdpE*), beta-lactams (*e.g., acrB, CRP, evgA*, *gadX, soxR*, *H-NS* etc.), carbapenems (*e.g., marA*, *soxS* etc.), cephalosporins (*e.g., acrA*), fluoroquinolones (*e.g., rsmA*, *emrA, emrR, emrB, mdtM, mdtH, mdtF, mdtE*), tetracyclines (*e.g., mdfA*, *emrY, emrK*), and vancomycin (e.g., *vanG*), representing a core set of resistance determinants shared between these strains (**Fig. 5**).

**Fig. 5.**
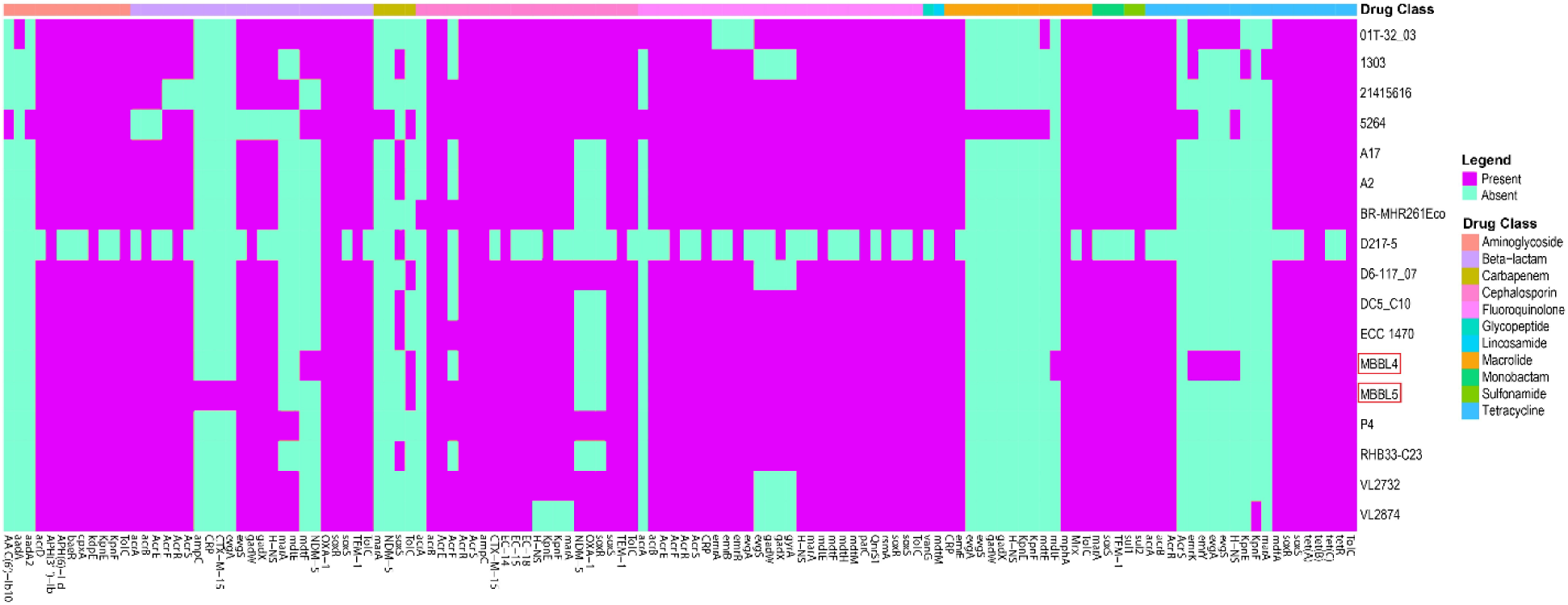
Heatmap of antibiotic resistance gene (ARG) profiles in 17 *E. coli* genomes. Salmon-colored boxes indicate the presence of ARGs, while mint blue-colored boxes indicate their absence. Antibiotic classes are color-coded in the legend as represented in the first row of the heatmap. Both MBBL4 and MBBL5 strains are highlighted in red boxes.

### Virulence gene repertoire and mechanistic insights in *E. coli* MBBL4 and MBBL5

To better elucidate the virulence potentials and mechanistic insights into pathogenesis of the two isolates, we conducted a comprehensive genomic analysis of their virulence repertoires. We found a significant variation in the composition of detected VFGs, indicating potential differences in pathogenicity between the genomes. The MBBL4 and MBBL5 genomes harbored 126 and 185 VFGs, respectively, facilitating virulence through mechanisms including adherence, effector delivery systems, siderophores, toxin production, invasion, regulatory proteins, bacterial growth/survival, and some uncharacterized functions (**Fig. 6, Table S7**). Adherence-related genes comprised the largest category in both strains, accounting for 76.2% of VFGs in MBBL4 and 73.5% in MBBL5. Within this category, both genomes encoded genes for flagella (42 in MBBL4 vs. 43 in MBBL5), fimbriae (13 vs. 26), pili (14 vs. 15), curli fibers (7 vs. 7), and other adhesion-related genes, including 4 general adhesion genes in MBBL4 and 12 adhesion-related genes (including 2 AIDA-I type adhesins) in MBBL5. Regarding secretion systems, MBBL4 contained Type II (2 genes) and Type III (14 genes), whereas MBBL5 carried a larger number of Type II (13 genes), Type III (17 genes), and additionally Type VI (3 genes) secretion system genes. Furthermore, both genomes harbor genes related to effector delivery, siderophores, toxin production, regulatory proteins, and bacterial growth/survival (**Fig. 6, Table S7**).

**Fig. 6.**
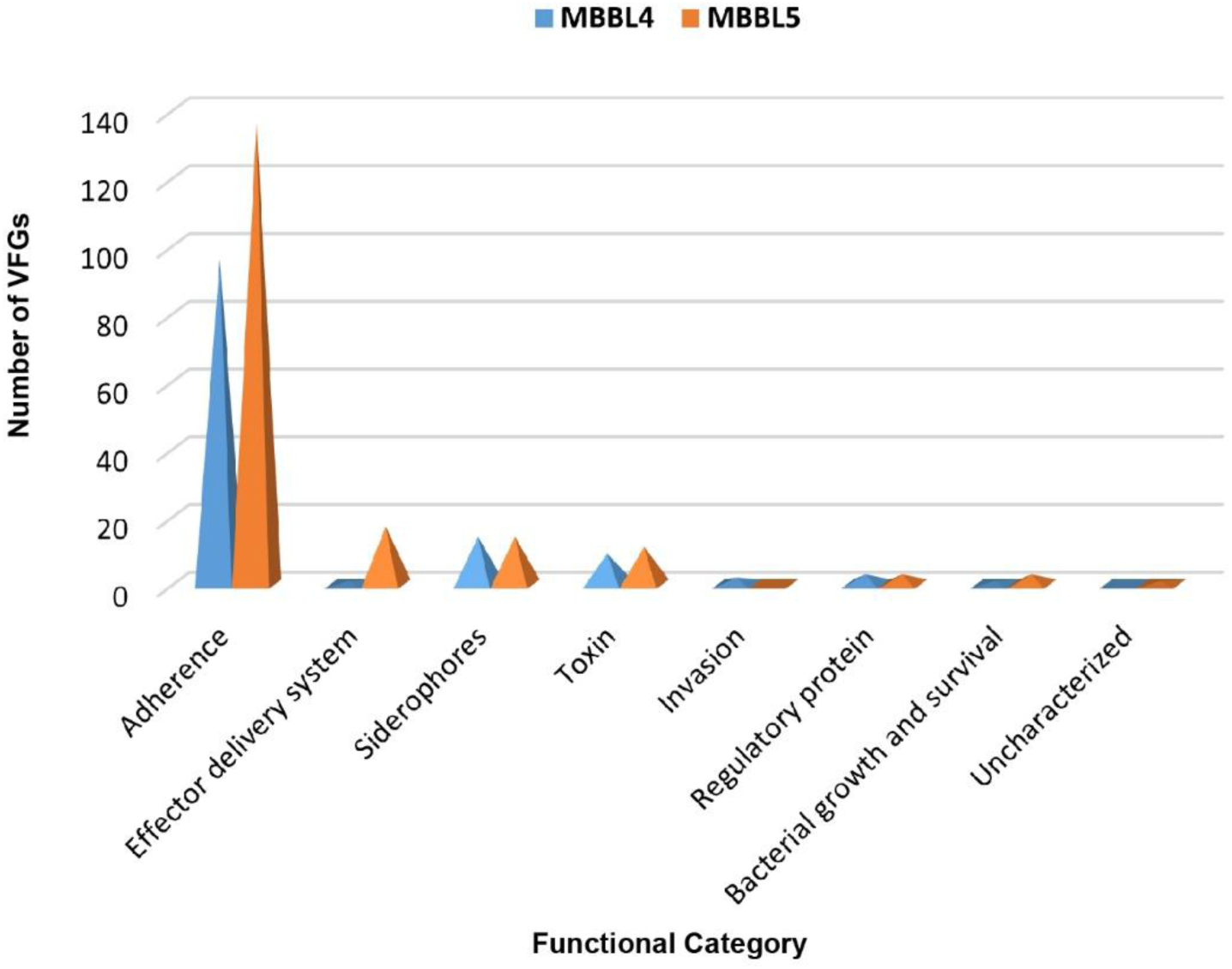
Overview of virulence functional potentials of *E. coli* MBBL4 and MBBL5.

### Genomic islands harboring resistance and virulence genes in *E. coli* MBBL4 and MBBL5

A key finding of this study was the identification of multiple GIs in both the MBBL4 and MBBL5 genomes (**Fig. 7**), which play a critical role in HGT. *E. coli* MBBL4 harbored 50 putative GIs (40 predicted by SIGI-HMM and 10 by IslandPath-DIMOB), with sizes ranging from 4,002 bp to 56,473 bp (**Fig. 7A**). Notably, six of these GIs were found to contain genes associated with VFGs or ARGs. The isolate encoded several curated ARGs, including *sul2* (WP_001043260.1), *evgA* (WP_000991370.1), and *ACHL6K_RS19350* (WP_000027057.1). Additionally, it harbored homologs of ARGs such as *ACHL6K_RS19340* (WP_000480968.1), *aph(3’’)-Ib* (WP_001082319.1), *emrY* (WP_001018731.1), *evgS* (WP_001355656.1). A pathogen-associated gene, *ACHL6K_RS08780* (WP_001198861.1), was also identified, along with a curated VFG, *ecpA* (WP_000730972.1) and a VFG homolog, *yfdV* (WP_000955028.1) (**Fig. 7A**). In contrast, *E. coli* MBBL5 contained 39 GIs (33 by SIGI-HMM and 6 by IslandPath-DIMOB), with lengths ranging from 4,157 bp to 56,409 bp (**Fig. 7B**). In this isolate, only three GIs contained genes associated with VFs or AMR. MBBL5 harbored a single curated resistance gene, *evgA*(WP_000991370.1). It also contained several homologs of ARGs, including *emrE* (WP_001070451.1), *emrY* (WP_039022714.1), *emrK*(WP_000435167.1), and *evgS* (WP_001307321.1). Additionally, one VFG homolog, *yfdV* (WP_000955028.1), was detected. No curated VFG or pathogen-associated genes were identified in MBBL5 (**Fig. 7B**). These genes are functionally linked to multidrug efflux, sulfonamide resistance, and adherence, contributing to the strain’s survival and pathogenic potential under antibiotic stress. In particular, *evgA* and *evgS* encode a two-component regulatory system that activates efflux pump expression, while *ecpA* encodes a major pilin subunit involved in adhesion to host cells. Furthermore, the higher number of GIs in MBBL4 compared to MBBL5 may be attributed to its exposure to a broader environmental microbial community or host-associated antibiotic stressors, which could enhance resistance and facilitate survival in diverse ecological niches.

**Fig. 7.**
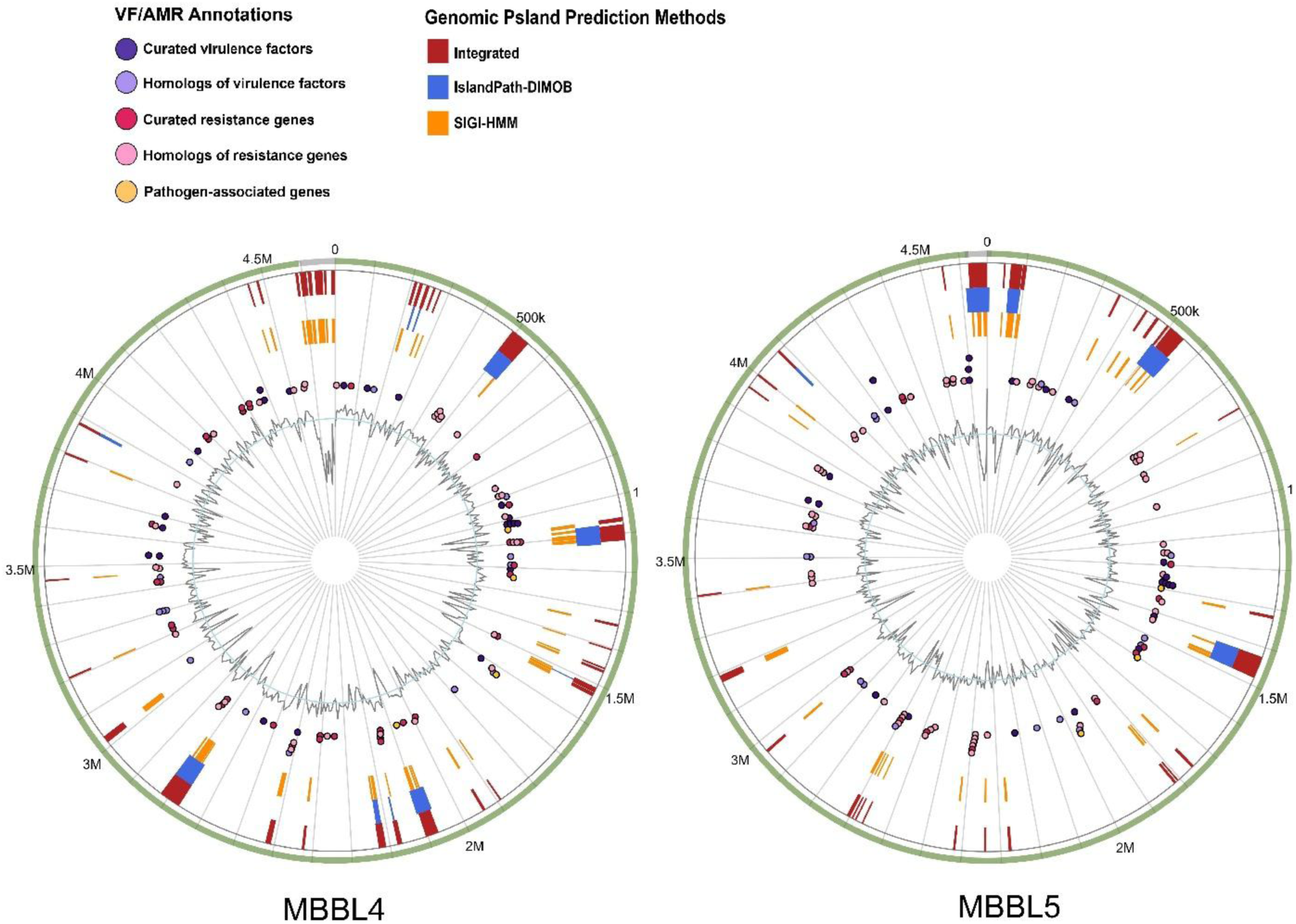
Comprehensive visualization of predicted genomic islands (GIs) and associated functional gene annotations in *E. coli* strains MBBL4 and MBBL5. Genomic islands predicted by the integrated method are shown in red, with additional predictions from IslandPath-DIMOB and SIGI-HMM displayed in blue and orange, respectively. Functional genes are color-coded as follows: curated virulence factor genes (VFGs) as purple circles, VFG homologs as lavender circles, curated antimicrobial resistance (AMR) genes as red circles, and pathogen-associated genes as golden yellow circles.

## Discussion

In this study, we performed high-throughput WGS of two MDR *E. coli* isolates, MBBL4 and MBBL5, isolated from bovine CM milk. Comprehensive genomic analysis of these isolates provides valuable insights into the genetic basis of AMR, virulence potential, and evolutionary dynamics of bovine mastitis-associated *E. coli*. The presence of diverse ARGs and VFGs in both genomes highlights the significant pathogenic potential of these isolates, which not only complicates mastitis treatment in dairy cattle but also raises concerns about possible zoonotic transmission primarily through consumption of unpasteurized milk or milk products and direct contact with infected animal or milk. Genomic features of MBBL4 and MBBL5 were consistent with previously reported mastitis-associated *E. coli* strains worldwide [14, 60, 61]. Phylogenetic analysis revealed that *E. coli* strains MBBL4 and MBBL5 belong to distinct evolutionary lineages. MBBL4 clustered with bovine mastitis-associated *E. coli* strains (P4, 1303, VL2732 and VL2874) [60–62] and the human bacteremia-associated *E. coli* strain 5264 (CP103540.1), suggesting potential zoonotic transmission risks. Conversely, MBBL5 aligned more closely with wildlife-derived fecal originated *E. coli* strains A2 and A17 [63], indicating possible environmental adaptation. ANI analysis also supported these relationships, with both strains showing over 96% identity to reference genomes. Pangenome analysis further demonstrated substantial genetic diversity, with MBBL4 and MBBL5 sharing 2,883 core genes but possessing distinct unique and accessory gene sets, emphasizing the adaptive potential of mastitis-associated *E. coli* [14, 59]. The identification of diverse MGEs, including plasmids, prophages, ISs, and RGPs, in both genomes underscores their extensive genetic diversity and adaptive capacity across various ecological niches [23, 64]. Although the circular plasmid (p0111) predicted in MBBL4 does not harbor any ARG or VFG, it could facilitate the horizontal transfer of these genes, thereby contributing to the global spread of MDR [65, 66]. Comparative analysis of ISs and prophages revealed notable differences between the isolates. MBBL5 exhibited a broader diversity and higher abundance of IS families, including ISNCY, IS3, IS21, and IS110, along with distinct prophage contents, which may influence HGT, virulence, and AMR, as prophages and IS elements are key drivers of genome diversification in pathogenic bacteria [67]. Interestingly, the close phylogenetic relationship of MBBL5 to wildlife-derived strains A2 and A17 suggests potential environmental transmission pathways, such as contamination of shared water sources or indirect contact between wildlife and agricultural environments. Despite its high nucleotide similarity to MBBL4, MBBL5 harbors unique MGEs, including prophages and plasmid-associated genes, which likely explain its distinct phylogenetic clustering and reflect functional divergence. Together, these findings highlight the role of wildlife and environmental reservoirs in shaping *E. coli* diversity and underscore the potential public health implications, including indirect risks of transmission through environmental contamination and the emergence of novel pathogenic strains.

Resistome analysis revealed that over 42% of ARGs were conserved across the studied *E. coli* strains, suggesting widespread intrinsic resistance mechanisms inherent to diverse *E. coli* lineages, irrespective of their isolation source, host, or geographical location. These findings are consistent with previous reports of high prevalence of resistance genes in *E. coli* from livestock in Bangladesh, particularly against beta-lactams, tetracyclines, and sulfonamides, highlighting the regional relevance of intrinsic ARGs [68, 69]. This widespread conservation of intrinsic ARGs in *E. coli* poses a significant threat, as it may limit treatment options and facilitate the persistence and spread of resistance across environments [70]. Although MBBL4 harbored higher number of ARGs (52) than MBBL5 (45), both strains shared core resistance determinants against a broad spectrum of antibiotics, including aminoglycosides, beta-lactams, carbapenems, cephalosporins, fluoroquinolones, glycopeptides, lincosamides, macrolides, monobactams, sulfonamides, and tetracyclines. This genomic resistome largely aligned with their observed phenotypic resistance profiles, with minor exceptions. The presence of conserved ARGs in *E. coli* isolates (MBBL4 and MBBL5) associated with bovine mastitis represents a critical challenge by severely restricting therapeutic options. This intrinsic and widespread resistance profile in *E. coli* genomes not only drives the persistence and dissemination of AMR within farm environments [14, 71] but also poses considerable zoonotic risks to humans through the consumption of contaminated milk and dairy products [71]. Addressing this growing threat necessitates the adoption of a comprehensive “One Health” strategy integrating animal, human, and environmental health. The comparative virulence gene analysis of *E. coli* MBBL4 and MBBL5 underscores significant pathogenic potential with strain-specific differences, indicating critical implications for animal and public health. In this study, the MBBL5 genome harboring a larger repertoire of VFGs (185) than MBBL4 (126), exhibits enhanced capabilities for colonization, immune evasion, and interbacterial competition. Both strains are highly enriched in adherence-related genes (> 70%), which are essential for biofilm formation, persistent colonization, and resistance to host defenses [72, 73]. Importantly, MBBL5 possesses additional secretion systems (Type VI) that contribute to delivering toxic effectors into host or competitor cells, intensifying its pathogenicity [74, 75]. These secretion systems may enhance the ability of MBBL5 to deliver effector proteins into mammary epithelial cells, exacerbating inflammation and tissue damage, and potentially contributing to more severe mastitis outcomes [76]. The presence of siderophore-mediated iron acquisition systems and multiple toxin genes in both strains underscores their potential to cause severe extraintestinal infections, including mastitis and zoonotic transmission [14, 77]. Many of these virulence features, including adhesins, siderophore systems, and secretion systems, are also commonly observed in well-characterized extra-intestinal pathogenic *E. coli* (ExPEC) strains, suggesting that MBBL4 and MBBL5 share pathogenic strategies with clinically important extraintestinal pathogens [78, 79]. However, MBBL5’s expanded VFG repertoire, particularly the presence of Type VI secretion system genes, may confer enhanced competitive and invasive capabilities relative to typical ExPEC strains. These findings highlight that MBBL5 may pose a greater threat due to its expanded virulence arsenal, facilitating more effective host invasion and persistence. In addition, the high pathogenicity scores, along with the large number of matched pathogenic gene families in both MBBL4 and MBBL5, reinforce their potential as zoonotic threats and highlight the clinical relevance of mastitis-associated *E. coli* in both animal and human health contexts. GIs play a crucial role in *E. coli* pathogenicity by facilitating HGT of virulence and resistance genes, enhancing adaptation, survival, host colonization, and disease-causing potential [80, 81]. The presence of multiple GIs in both *E. coli* genomes highlighted their key role in HGT and genome evolution. In MBBL4, GIs carried several important ARGs (e.g., *sul2*, *aph(3’’)-Ib*) and a VFG (*ecpA*), indicating strong resistance and virulence potential. Although MBBL5 had fewer GIs, they still contained ARGs (e.g., *evgA*, *emrE*, *emrY*) and a VFG homolog (*yfdV*), showing its ability to adapt in diverse environment. Additionally, several genes within the GIs may contribute to environmental persistence, enabling the bacteria to survive outside the host and contribute to the transmission within the farm environment. These findings underscore the pivotal role of GIs in driving genomic plasticity, shaping the evolution of resistance and virulence, and contributing to the emergence of high-risk *E. coli* lineages. Importantly, they emphasize the need for continuous longitudinal genomic surveillance to detect and monitor MGEs that may facilitate the spread of clinically relevant traits across populations and ecosystems. Although this study offers a comprehensive genomic characterization of two *E. coli* strains (MBBL4 and MBBL5), a notable limitation lies in the small number of genomes analyzed (n = 2), which constrains the generalizability of the findings to the wider *E. coli* population associated with bovine mastitis. A higher number of samplings would provide a more comprehensive understanding of *E. coli* diversity and genomic features. While the genomic data yielded valuable insights into potential virulence and resistance traits, the study does not establish a causal link between these genetic features and the corresponding phenotypic traits such as biofilm formation and in vivo virulence. Additionally, the absence of *in vivo* validation limits the translational relevance of these findings for clinical application.

## Conclusion

This study provides comprehensive genomic insights into two MDR *E. coli* isolates from bovine CM milk, revealing the intricate relationship between AMR, virulence, and MGEs. The coexistence of diverse resistance and virulence determinants, along with abundant ISs, prophages, and GIs, highlights their remarkable adaptability and potential for cross-species transmission. These findings underscore the growing threat posed by mastitis-associated *E. coli* as reservoirs of MDR within the dairy environment. Consequently, this study underscores the critical necessity of integrating genomic surveillance into a One Health framework. This study provides a scientific basis for policymakers to mandate the development of National Mastitis Control Guidelines, integrate AMR surveillance into the existing livestock extension services, and promote biosecurity protocols and rational antibiotic use on dairy farms. Integrating WGS into One Health surveillance frameworks is essential to monitor genomic evolution, identify high-risk lineages, and guide evidence-based interventions. Proactive monitoring of such clonal lineages is paramount for developing targeted interventions to mitigate their spread and safeguard both veterinary and public health.

## Supporting information

Supplementary Tables S3-S5.docx

## Declarations

### Ethical approval

The Animal Research Ethics Committee (AREC) of the Gazipur Agricultural University (Gazipur, Bangladesh) reviewed and approved the experimental procedures of this study (reference number FVMAS/AREC/2023/6679 [16 January 2023]). Written informed consent was obtained from each participating dairy farmer prior to the inclusion of their animals in this study, ensuring ethical compliance and voluntary participation.

### Clinical trial number

Not applicable.

### Consent for publication

Not applicable.

### Authors contributions

M.N.H. conceived and designed the study. N.S., K.Y.A, M.A.A.G. and M.M.R. performed experiment, curated, analyzed and visualized data, and wrote original draft. Z.C.D., T.I. and M.N.H. critically reviewed and edited the manuscript. The final manuscript was read and approved by all authors.

### Data availability

The *16S rRNA* gene sequences of *E. coli* strains MBBL4 and MBBL5 have been deposited in NCBI GenBank under accession numbers [PV855738](https://submit.ncbi.nlm.nih.gov/subs/genbank/SUB15426323/overview) and [PV855739](https://submit.ncbi.nlm.nih.gov/subs/genbank/SUB15426343/overview), respectively. The whole genome shotgun sequences for both strains are available in GenBank and the NCBI Sequence Read Archive (SRA) under BioProject accession PRJNA1171635. The genome versions reported in this study are [JBINJR000000000.1](https://www.ncbi.nlm.nih.gov/nuccore/JBINJR000000000) for MBBL4 and [JBINJQ000000000.1](https://www.ncbi.nlm.nih.gov/nuccore/JBINJQ000000000) for MBBL5.

