## Supplementary Tables S1, S2, S6 and S8 for "Genomic insights into multidrug resistant *Escherichia coli* from bovine mastitis in Bangladesh"

**Table S1**. List of assembly ID, name of the strains, source and host of the *E. coli* strains.

| **Sl. No.** | **Assembly ID** | **Strain** | **Source** | **Country** | **Host** |
| --- | --- | --- | --- | --- | --- |
| 1 | GCA_043952105.1 | MBBL4 | Bovine mastitic milk | Bangladesh | Bovine |
| 2 | GCA_043952135.1 | MBBL5 | Bovine mastitic milk | Bangladesh |  |
| 3 | [GCA_000259425.1](https://www.ncbi.nlm.nih.gov/datasets/genome/GCA_000259425.1/) | P4 | Bovine mastitic milk | UK |  |
| 4 | [GCA_024623645.1](https://www.ncbi.nlm.nih.gov/datasets/genome/GCA_024623645.1/) | 01T-32/03 | Bovine mastitic milk | Brazil |  |
| 5 | [GCA_024623585.1](https://www.ncbi.nlm.nih.gov/datasets/genome/GCA_024623585.1/) | 23T-166/22 | Bovine mastitic milk | Brazil |  |
| 6 | [GCA_030360205.1](https://www.ncbi.nlm.nih.gov/datasets/genome/GCA_030360205.1/) | MAHK_SCM_BAU_30A | Bovine mastitic milk | Bangladesh |  |
| 7 | [GCA_000829985.1](https://www.ncbi.nlm.nih.gov/datasets/genome/GCA_000829985.1/) | 1303 | Bovine mastitic milk | Germany |  |
| 8 | [GCA_000831565.1](https://www.ncbi.nlm.nih.gov/datasets/genome/GCA_000831565.1/) | ECC-1470 | Bovine mastitic milk | USA |  |
| 9 | [GCA_022631305.1](https://www.ncbi.nlm.nih.gov/datasets/genome/GCA_022631305.1/) | BR-MHR268Eco | Bovine mastitic milk | Bangladesh |  |
| 10 | [GCA_022631295.1](https://www.ncbi.nlm.nih.gov/datasets/genome/GCA_022631295.1/) | BR-MHR261Eco | Bovine mastitic milk | Bangladesh |  |
| 11 | [GCA_021045155.1](https://www.ncbi.nlm.nih.gov/datasets/genome/GCA_021045155.1/) | MEZEC95 | Bovine mastitic milk | South Africa |  |
| 12 | [GCA_001441325.1](https://www.ncbi.nlm.nih.gov/datasets/genome/GCA_001441325.1/) | VL2732 | Bovine mastitic milk | Israel |  |
| 13 | [GCA_001441335.1](https://www.ncbi.nlm.nih.gov/datasets/genome/GCA_001441335.1/) | VL2874 | Bovine mastitic milk | Israel |  |
| 14 | [GCA_000731355.1](https://www.ncbi.nlm.nih.gov/datasets/genome/GCA_000731355.1/) | [D6-113.11](https://www.ncbi.nlm.nih.gov/taxonomy/1400022) | Bovine mastitic milk | France |  |
| 15 | [GCA_000731455.1](https://www.ncbi.nlm.nih.gov/datasets/genome/GCA_000731455.1/) | D6-117.07 | Bovine mastitic milk | France |  |
| 16 | [GCA_016113265.1](https://www.ncbi.nlm.nih.gov/datasets/genome/GCA_016113265.1/) | 22710987 | Bovine mastitic milk | Canada |  |
| 17 | [GCA_016113085.1](https://www.ncbi.nlm.nih.gov/datasets/genome/GCA_016113085.1/) | 21415616 | Bovine mastitic milk | Canada |  |
| 18 | [GCA_024917815.1](https://www.ncbi.nlm.nih.gov/datasets/genome/GCA_024917815.1/) | 5264 | Human bacteremia | USA | Human |
| 19 | [GCA_024918155.1](https://www.ncbi.nlm.nih.gov/datasets/genome/GCA_024918155.1/) | 4069 | Human bacteremia | USA |  |
| 20 | [GCA_024584925.1](https://www.ncbi.nlm.nih.gov/datasets/genome/GCA_024584925.1/) | D217-5 | Human (child) diarrhoea | Thailand |  |
| 21 | [GCA_024584885.1](https://www.ncbi.nlm.nih.gov/datasets/genome/GCA_024584885.1/) | 0-4 | Human (adult) diarrhoea | Brazil |  |
| 22 | [GCA_028898885.1](https://www.ncbi.nlm.nih.gov/datasets/genome/GCA_028898885.1/) | NB4833 | Human (urine) cervical cancer | China |  |
| 23 | [GCA_027944795.1](https://www.ncbi.nlm.nih.gov/datasets/genome/GCA_027944795.1/) | DETEC-P666 | ICU patient oral swab | China |  |
| 24 | [GCA_022493855.1](https://www.ncbi.nlm.nih.gov/datasets/genome/GCA_022493855.1/) | A2 | Platypus (feces) | Australia | Other animals |
| 25 | [GCA_022493555.1](https://www.ncbi.nlm.nih.gov/datasets/genome/GCA_022493555.1/) | A17 | Otter (feces) | Mexico |  |
| 26 | [GCA_030013845.1](https://www.ncbi.nlm.nih.gov/datasets/genome/GCA_030013845.1/) | ETEC1721 | Pig feces (enteric colibacillosis) | Belgium |  |
| 27 | [GCA_013823705.1](https://www.ncbi.nlm.nih.gov/datasets/genome/GCA_013823705.1/) | RHB33-C23 | Pooled sheep fecal samples | UK |  |
| 28 | [GCA_014104555.1](https://www.ncbi.nlm.nih.gov/datasets/genome/GCA_014104555.1/) | RHB37-C11 | Pooled sheep fecal samples | UK |  |
| 29 | [GCA_023369735.1](https://www.ncbi.nlm.nih.gov/datasets/genome/GCA_023369735.1/) | DC5_C10 | Water | USA | Environmental |
| 30 | [GCA_029717685.1](https://www.ncbi.nlm.nih.gov/datasets/genome/GCA_029717685.1/) | RHB07-SO-C04 | Soil | UK |  |

**Table S2.** List of core, accessory and strain specific genes in 17 *E. coli* genomes.

| Genome ID | Core genes | Accessory genes | Strain specific genes |
| --- | --- | --- | --- |
| 01T-32_03 | 2883 | 392 | 22 |
| 1303 | 2883 | 538 | 29 |
| 21415616 | 2883 | 473 | 19 |
| 5264 | 2883 | 429 | 43 |
| A17 | 2883 | 477 | 10 |
| A2 | 2883 | 550 | 31 |
| BR-MHR261Eco | 2883 | 509 | 23 |
| D217-5 | 2883 | 516 | 5 |
| D6-117_07.11 | 2883 | 483 | 46 |
| DC5_C10 | 2883 | 495 | 20 |
| ECC-1470 | 2883 | 523 | 20 |
| MBBL4 | 2883 | 380 | 21 |
| MBBL5 | 2883 | 468 | 9 |
| P4 | 2883 | 505 | 35 |
| RHB33-C23 | 2883 | 509 | 4 |
| VL2732 | 2883 | 457 | 31 |
| VL2874 | 2883 | 488 | 33 |

**Table S6.** Summary of plasmid sizes in 17 *E. coli* genomes.

| Genome ID | Total_Plasmid_Size (bp) |
| --- | --- |
| VL2732 | 0 |
| VL2874 | 0 |
| A2 | 0 |
| A17 | 0 |
| MBBL4 | 39271 |
| MBBL5 | 0 |
| D217-5 | 0 |
| RHB33-C23 | 0 |
| 5264 | 10272 |
| 21415616 | 0 |
| P4 | 17124 |
| 01T-32_03 | 12499 |
| BR-MHR261Eco | 12174 |
| ECC-1470 | 100061 |
| DC5_C10 | 0 |
| 1303 | 99630 |
| D6-117_07.11 | 0 |

**Table S7:** Functional annotation of virulence factor genes (VFGs) and their virulence mechanisms in *E. coli* strains MBBL4 (n = 126) and MBBL5 (n = 185).

| **Sl. No.** | **Functional category** | **Sub-category** | **MBBL4** | **MBBL5** |
| --- | --- | --- | --- | --- |
| 1 | Adherence | Adhesion | 4 (Z1307, b2854, eaeH, and cadA) | 10 (cadA, eaeH, b2854, Z0263/0265/1307/2200/2201/2204/2206) |
|  |  | Flagella | 42 (flgA/B/C/D/E, flhA/B/C/D/E, fliA/D/E/F/G, flk, motA/B, ycfz, etc.) | 43 (flhA/B/C/D/E, flgA/B/C/D/E, fliA/C/D/E/F/G/H, flk, motA/B, ycfz, etc.) |
|  |  | Fimbriae | 13 (ycbF/Q/R/S/T/V, fimA/C/D/F/G/H/I) | 26 (cfaA/B/C/D, fimA/B/C/D/E/F/G/H/I, matF, etc.) |
|  |  | Curli fibers | 7 (csgA/B/C/D/E/F/G) | 7 (csgA/B/C/D/E/F/G) |
|  |  | Pili | 14 (ppda/b/c/D, ygdb, yggr, hofq/C, ecpA/B/C/D/E, and ecpR) | 15 (ecpA/B/C/D/R, hofB/C/q, ppda/b/c/D, ycbU, ygdb, and yggr) |
|  |  | AIDA-I type | -- | 2 (ehaA and ehaB) |
|  |  | Type II secretion system | 2 (gspM and gspo) | 13 (b2972, gspC/D/E/F/G/H/I/J/K/L/M, and yghg) |
|  |  | Type III secretion system | 14 (eprH/I/J/K/O/P/Q/R/S, espL3/L4/X4/Y1, and ygeH) | 17 (epaO/P/Q/R/S, eprH/I/J/K, espL1/L3/L4/R1/X1/X4/X5, and ygeH) |
|  |  | Type IV secretion system | -- | 3 (clpV, hcp, and vgrG) |
| 2 | Effector delivery system | ACE T6SS, Effector delivery system | -- | 13 (aec17, aec18, aec19, aec22, aec23, aec24, aec25, aec26, aec28, aec29, aec30, aec31, and aec32) |
|  |  | ETT2, Effector delivery system | 1 (etrA) | 4 (ECs3712, ECS88_3547, etrA, and ygeG) |
| 3 | Siderophore producing |  | 14 (entA, entB, entC, entD, entE, entF, entS, fepA, fepB, fepC, fepD, fepE, fepG, and fes) | 14 (entA, entB, entC, entD, entE, entF, entS, fepA, fepB, fepC, fepD, fepE, fepG, and fes ) |
| 4 | Toxin producing |  | 9 (cheA, cheB, cheR, cheW, cheY, cheZ, tar/cheM, hlyE, and ibeB) | 11 (cheA, cheB, cheR, cheW, cheY, cheZ, hlyE, ibeB, ibeC, iss2, and tar/cheM) |
| 5 | Invasion |  | 2 (aslA and ompt) | -- |
| 6 | Regulatory protein |  | 3 (natb, int, and gadX) | 3 (nada, nadb, and gadX) |
| 7 | Bacterial growth and survival |  | 1 (artj) | 3 (artj, orgA and orgB) |
| 8 | Uncharacterized |  | -- | 1 (UMNK88_238) |
